# Combining molecular gut content analysis and functional response models shows how body size affects prey choice in soil predators

**DOI:** 10.1101/113944

**Authors:** Bernhard Eitzinger, Björn C. Rall, Michael Traugott, Stefan Scheu

## Abstract

1. Predator-prey interactions are a core concept of animal ecology and functional response models provide a powerful tool to predict the strength of trophic links and assess motives for prey choice. However, due to their reductionist set-up, these models may not display field conditions, possibly leading to skewed results.
2. We tested the validity of functional response models for multiple prey by comparing them with empirical data from DNA-based molecular gut content analysis of two abundant and widespread macrofauna soil predators, lithobiid and geophilomorph centipedes.
3. We collected soil and litter dwelling centipedes, screened their gut contents for DNA of nine abundant decomposer and intraguild prey using specific primers and tested for different prey and predator traits explaining prey choice. In order to calculate the functional response of same predators, we used natural prey abundances and functional response parameters from published experiments and compared both approaches.
4. Molecular gut content results showed that prey choice of centipedes is driven by predator body size and prey identity. Results of functional response models significantly correlated with results from molecular gut content analysis for the majority of prey species.
5. Overall, the results suggest that functional response models are a powerful tool to predict trophic interactions in soil, however, species-specific traits have to be taken into account to improve predictions.

## Introduction

Analysis of consumer-resource interactions is key to understand the structure and dynamics of food webs, eventually explaining composition, stability and development of communities and ecological processes coupled with them. Depending on the specific problem and scale of feeding interactions, 21^st^ century ecologists are in the comfortable position to select from a broad spectrum of methods, from field observations to molecular tracking of nutrients and DNA in the consumer’s body. Measuring the functional response, i.e. the intake rate of a consumer (hereafter referred to as *predator*) as a function of food resource (hereafter referred to as *prey*) density has been demonstrated to be a powerful method not only to track feeding interactions but also to assess the interaction strength (Holling 1959). Based on a small set of parameters including densities and body sizes of prey and predator, functional response models allow predicting general patterns and mechanisms of trophic interactions in very different systems, spanning from *Daphnia* water fleas feeding on phytoplankton to wolf packs preying on moose (Sarnelle & Wilson 2008, Messier 1994). The approach allows investigating feeding interactions on a large scale, and can be modified to include changes in body size (Hansen *et al.* 1997, Pawar *et al.* 2012, Rall *et al.* 2012), ambient temperature (Hansen *et al.* 1997, Englund *et al.* 2011, Rall *et al.* 2012) as well as habitat structure (Hauzy *et al.* 2010, Kalinkat *et al.* 2013a, Kalinkat & Rall 2015).

The simplicity of functional responses, however, may come at the cost of accuracy. Functional response curves, in particular those of invertebrate species, are typically based on single-prey-predator laboratory feeding trials, which lack many characteristics of natural settings. Among these potentially important characteristics are habitat structure, presence of competitors and alternative prey as well as different physiological states of prey and predator (e.g. sick prey). Thus, functional response models based on idealized laboratory settings may be of limited use to predict feeding interactions in the field. Here, other methods apply, which allow us to analyse the character and intensity of predator-prey interactions under natural settings directly in the field.

DNA-based molecular gut content analysis offers a state of-the-art technique (Pompanon *et al.* 2012; Traugott *et al.* 2013) to identify trophic links under challenging conditions, from sea shores (Peters *et al.* 2014), over arctic tundra (Wirta *et al.* 2015) to arable soils (Wallinger *et al.* 2014). Using specifically designed PCR assays targeting prey DNA in a predator’s gut, species-specific trophic interactions can be tracked even several days after the feeding event, allowing unravelling of trophic links in unprecedented detail (Eitzinger *et al.* 2013). Hence, molecular gut content analysis allows to empirically assess complex trophic interactions in the field and provides the opportunity to evaluate functional response models under natural conditions.

We adopted this approach for the first time using for a soil predator-prey system in European deciduous forests. Here, we analysed the predation frequency on extra- and intraguild prey of centipedes (Chilopoda, Myriapoda), widespread generalist predators in the litter and soil layers of temperate forests (Lewis 1981; Poser 1988) using predictive models from functional response experiments and compare these with empirically quantified trophic links using molecular gut content analysis. By the combined use of both approaches we aimed at achieving an integrated view of food web interactions in complex systems and evaluate the suitability and effectiveness of the approaches for analysing trophic interactions.

Centipedes, in particular lithobiid (Lithobiidae) and geophilomorph (Geophilomorpha) species prey on a variety of prey taxa including Collembola, Diptera larvae and Lumbricidae (Günther *et al.* 2014). Lithobiids predominantly colonize the litter layer and perform a sit-and-wait strategy of prey capture, whereas geophilomorph centipedes are active hunters in crevices of the mineral soil (Lewis 1981; Poser 1988; Eitzinger *et al.* 2013). Prey capture of centipedes specifically depends on body size indicating an allometric relationship between predator and prey size (Schneider *et al.* 2012, Günther *et al.* 2014). Typically, small predators have narrow diets while large predators feed on a wider range of prey including higher trophic level taxa, i.e. intraguild prey (Woodward & Hildrew 2002; Riede *et al.* 2011). Body-size dependent prey-switching, coupled with feeding on intraguild prey may be a key factor reducing dietary niche overlap (Woodward & Hildrew 2002). Moreover, this might explain coexistence of different centipede species and other predators in forest soils. Studies employing functional response models suggested that the body size acts as a supertrait, explaining most of the variance in predator-prey interactions in soil systems (Vucic-Pestic *et al.* 2010, Kalinkat *et al.* 2013b). Hence, allometry-based functional response models may be applied to many different predator-prey-interactions.

Based on the generalised allometric functional response model by Kalinkat *et al.* (2013b), we calculated body-size dependent trophic interaction strength of centipede predators as a function of natural abundances of different prey groups present in soil of unmanaged beech forests in central Germany. We then analysed the gut content of field-collected centipedes from the same forests using nine group- and five species-specific primers for DNA of extra- and intraguild prey taxa. We hypothesized that (i) feeding interactions of centipedes are driven by predator-prey body-size ratios rather than by taxonomy, and that (ii) functional response models can correctly predict actual feeding interactions in a complex system such as soil.

## Materials and Methods

### Sampling

Invertebrate predators were collected in four unmanaged beech forests (> 120 years old) within the national park Hainich near Mülverstedt (Thuringia, Germany). Each study plot spanned 100 × 100 m and formed part of the Biodiversity Exploratories, an integrated biodiversity project (Fischer *et al.* 2010). To investigate trophic links during periods of maximum invertebrate activity, we sampled animals in autumn and spring/early summer, each represented by four sampling dates (October 8, 20 and 28 and November 3, 2009; June 15, 24 and 29 and July 8, 2010). Centipedes were collected by sieving litter, transferred individually to 1.5 mL microcentrifuge tubes and placed immediately at -20 °C.

To record the species spectrum and abundance of prey organisms, two large (20 cm diameter, 10 cm deep) and two small (5 cm diameter, 10 cm deep) soil cores per plot were taken in May of 2008 and 2011 (Klarner *et al.* 2014). Animals were extracted using a high gradient extractor (Kempson *et al.* 1963), stored in 75% ethanol and identified to the species level (except dipteran larvae). Additionally, lumbricids were collected by hand after application of mustard solution (Eisenhauer *et al.* 2008). Average densities between the two sampling dates were taken to represent prey density at the sampling dates of centipedes. We assume this to be justified as soil arthropod composition and density changes little between years (Bengtsson 1994).

A total of 532 field-caught *Lithobius* spp. and 65 geophilomorph centipedes were identified to species level using the keys of Eason (1964) and Latzel (1880). Further, we determined developmental stages and body length of each individual. Body mass of lithobiid centipedes was calculated using the following equation:

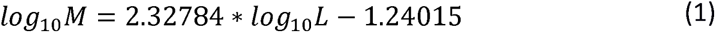

where *M* is the fresh body mass and *L* the body length of individuals. The equation is based on body length - body mass relationship of 560 lithobiid individuals used in laboratory studies (Eitzinger *et al.* 2014). Based on body size of collected specimens from the study site the body mass of geophilomorph centipedes and all prey taxa was calculated using formulas given in Gowing and Recher (1984) and Mercer (2001). Body mass (for predator and prey) and prey abundance were log_10_-transformed prior to statistical analyses.

### DNA extraction

Centipedes were subjected to CTAB-based DNA-extraction protocol (Juen & Traugott 2005) with modifications given in Eitzinger *et al.* (2013). DNA extracts were purified using Geneclean Kit (MP Biomedicals, Solon, OH, USA). To test for DNA carry-over contamination a blank control was included within a batch of 47 individuals. None was found when testing all extracts for false negatives and false positives, using the universal invertebrate primer pair LCO1490/HCO2198 (Folmer *et al.* 1994) amplifying a *c*. 700 bp fragment of the cytochrome *c* oxidase subunit I gene (*COI*). Each 10 μL PCR contained 5 μL PCR SuperHot Mastermix (2×), 1.25 mM MgCl_2_ (both Geneaxxon, Ulm, Germany), 0.5 μL bovine serum albumin (BSA, 3%; Roth, Karlsruhe, Germany), 0.5 μM of each primer and 3 μL of DNA extract. PCR cycling conditions were 95 °C for 10 min followed by 35 cycles at 95 °C for 30 s, 48 °C for 30 s, 72 °C for 90 s and a final elongation at 72 °C for 10 min. PCR products were separated in 1% ethidium bromide-stained agarose gels and visualized under UV-light.

### Screening predators for prey DNA

DNA extracts were screened for five extraguild and three intraguild prey (i.e. other predators) taxa using group-specific primers. PCR mixes and thermocycling conditions were the same as above only differing in applied primers, an elongation step at 72 °C for 45 s and the primer pair-specific annealing temperature. Geophilomorph centipedes additionally were tested for consumption of *Lithobius* spp. intraguild prey. All predator samples scoring positive for Collembola were subsequently tested for abundant Collembola species *Ceratophysella denticulata, Folsomia quadrioculata, Lepidocyrtus lanuginosus, Protaphorura armata and Pogonognathellus longicornis* (for primers and annealing temperature see Table S1, Supporting Information).

Specificity of the PCR assays was warranted by testing against a set of up to 119 non-target organisms (Eitzinger *et al.* 2013). PCR products were separated using the capillary electrophoresis system QIAxcel (Qiagen, Hilden, Germany); fragments of the expected size and a relative fluorescent value ≥ 0.1 RFU were scored as positive. PCR products showing no result were retested once.

### Statistical analysis

To compare prey DNA detection rates between predator taxa at the *P* < 0.05 level, 95% tilting confidence intervals (CI; Hesterberg *et al.* 2003) were calculated by 9999 bootstrap resamples using s-plus 8.0 (Insightful Corporations, Seattle, WA, USA).

Relationships between prey detection rates and predator identity, predator body mass, square of predator body mass, predator development stage (immature or adult), prey identity, prey body mass and prey abundance were analysed by generalized linear models (GLM) in R 2.12.2 (R Development Core Team 2011) using the function glm {stats}. Based on Akaike information criterion (AIC) we selected the most parsimonious model (Burnham and Anderson 2004). Prey DNA detection data was coded as binary (prey DNA present or absent).

A multi-prey functional response model was used to calculate feeding rates *F* of centipede predator *i* and prey *j* when alternative prey organisms *k* are present (note that *k* includes j; Kalinkat *et al.* 2011):

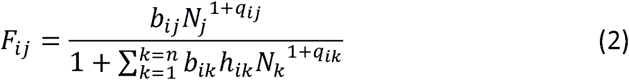

with *N* the prey density (individuals/m^2^), *n* the number of alternative prey items, *h* [s] the handling time (time for killing, ingesting and digesting prey), *b* the capture coefficient and *q* the scaling exponent that converts hyperbolic type-II (q = 0) into sigmoid type-III (q> 0) functional responses (Kalinkat *et al.* 2013b). We used prey-specific body masses [g] and values for generalised allometric functional response (Kalinkat *et al.* 2013b) to calculate b, *h* and *q* for each of the eight most important prey groups and added plot-specific prey density data (see above). The relative proportion of each of the eight prey-specific feeding rates per plot and for all plots combined was measured, resulting in prey-specific feeding ratios, *Frel:*

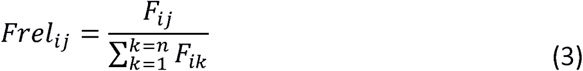

Additionally, we related both prey detection and feeding ratios to body size of predators.

For each prey group, we then compared the relative proportion of prey in the predator’s diet with the proportion of prey-DNA-positive predators using Pearson’s correlation coefficient in R 2.12.2.

## Results

### Centipede community

Among the 597 centipedes collected during the sampling periods, nine species of lithobiid (*Lithobius aulacopus, L. crassipes, L. curtipes*, *L. dentatus, L. melanops, L. muticus, L. mutabilis, L. nodulipes* and *L. piceus)* and three species of geophilomorph centipedes *(Geophilus* sp., *Schendyla nemorensis, Strigamia acuminata)* of both sexes and different developmental stages were identified. Body sizes / body masses ranged between 2-18 mm / 0.28 - 48.07 mg in lithobiids and 8-47 mm / 1.58 - 16.70 mg in geophilomorph centipedes.

### Prey DNA screening

A total of 532 *Lithobius* spp. and 65 geophilomorph centipedes collected at the eight sampling dates were tested for DNA of five and four extra- and intraguild prey taxa, respectively. Per sampling date 41-91 *Lithobius* spp. and 4-12 geophilomorph centipedes were investigated.

DNA of each of the prey organisms tested could be detected in at least one predator individual. Lithobiid predators were significantly more often tested positive for Collembola than for any other prey group (Fig. 1A). Detection rates of Diptera and Lumbricidae were significantly higher than those of other extraguild prey, such as Isopoda and Oribatida. Intraguild prey formed only a minor fraction of lithobiid prey: detection frequencies of Mesostigmata were followed by Staphylinidae and Araneida. In 69 predators two or three prey taxa were detected in one individual. The lithobiids which tested positive with the general Collembola primers (*n*=141) consumed significantly more *Folsomia quadrioculata* than any other of the four tested Collembola species (Fig. 1B).

**Fig 1.**
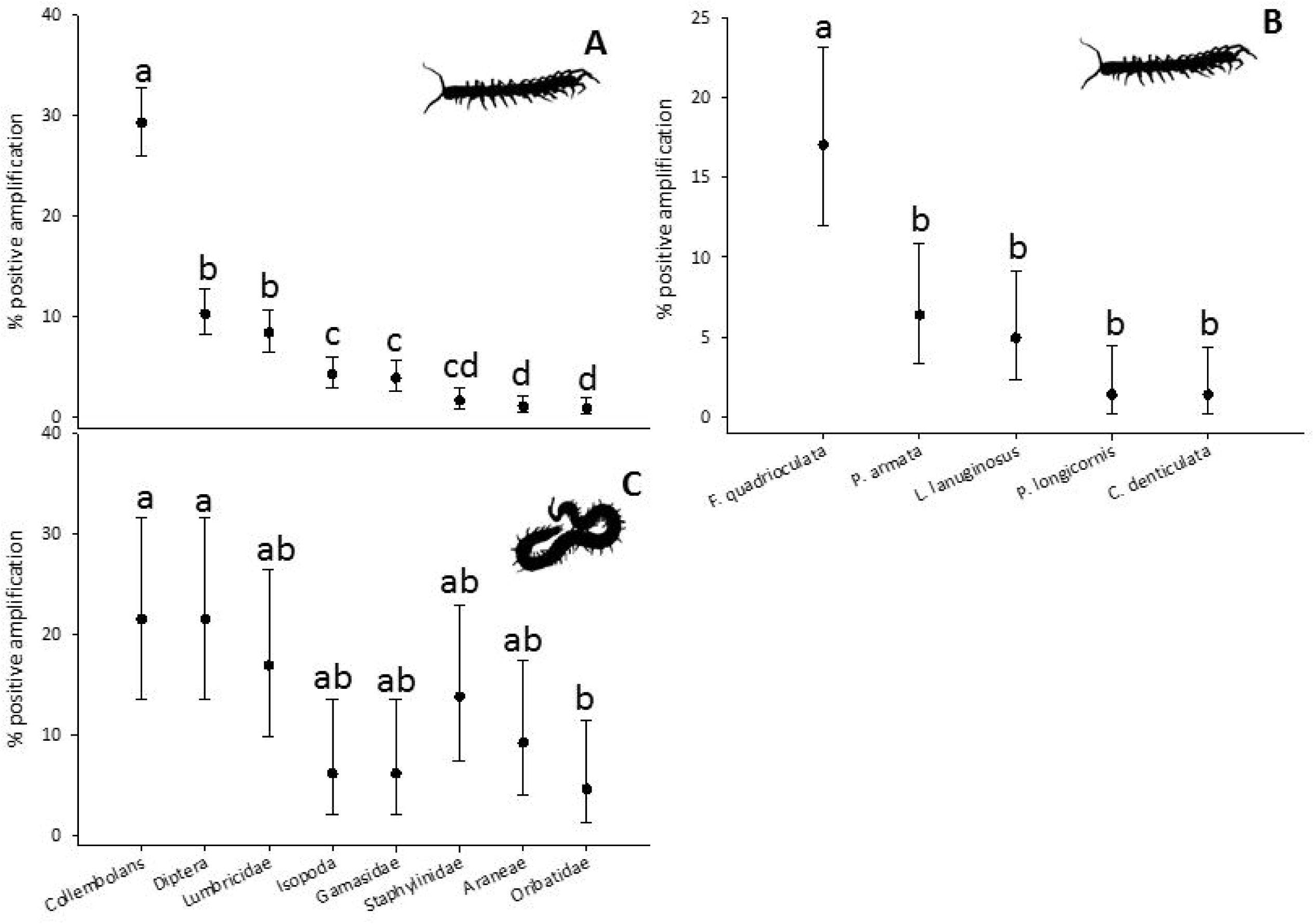
Prey detection rates of lithobiid (A; n= 532) and geophilomorph centipedes (C; n=65) sampled in autumn 2009 and spring 2010. Specimens tested positive for Collembola prey (B; n=141) further were tested for Collembola prey species. Error bars indicated 95% confidence intervals and letters denote significant differences in DNA detection rates at P < 0.05.

In geophilomorph centipedes extraguild prey, such as Collembola and Diptera, were most often detected followed by Lumbricidae, Isopoda and Oribatida (Fig. 1C). Detection rates for intraguild prey were highest for Staphylinidae, followed by Araneida and Mesostigmata. None of the five Collembola species could be detected in geophilomorph centipedes tested positive for Collembola. In 14 geophilomorph centipedes two or three prey taxa were detected simultaneously.

### Factors influencing prey consumption

We selected the most parsimonious model based on AIC comparison, thereby rejecting models containing factors centipede identity and development stage. Overall, lithobiid feeding was significantly affected by prey identity and predator body mass (Table 1), with preferences of predators for certain prey sizes. For Collembola and Lumbricidae prey, the probability of prey detection in relation to predator body mass followed a unimodal curve, peaking at body masses of 6.3 mg and 4.9 mg, respectively (Fig. 2). In contrast, detection probability of Diptera prey increased exponentially with predator body mass, indicating that Diptera are increasingly fed on by larger lithobiids while being rejected by smaller ones. Prey detection probabilities for Oribatida, Mesostigmata, Staphylinidae and Isopoda, despite being generally low, also increased with predator body mass, with the curve flattening at 25, 60, 62 and 69 mg predator body mass, respectively. Feeding on another intraguild prey, Araneida, however, showed a steady decrease with body mass. Feeding of geophilomorph centipedes varied with prey identity, predator body mass (including square of predator body mass) and prey abundance (Table S2, Supporting Information). In contrast to lithobiids, detection rates followed a unimodal curve for each of the prey taxa (Fig. S3, Supporting Information).

**Table 1.** Results of Generalized linear model (GLM) on the effect of predator body mass, square of predator body mass, prey identity and the two-way interactions on the detection of prey DNA in *Lithobius* predators. Significant effects are highlighted in bold. Df: degrees of freedom

| Variable | Df | Deviance | Resid. Df | Resid. Dev | P(>\|Chi\|) |
| --- | --- | --- | --- | --- | --- |
| NULL |  |  | 4247 | 2270.2 |  |
| Log<sub>10</sub> predator body mass | 1 | 5.38 | 4246 | 2264.8 | **0.0204** |
| Prey identity | 7 | 386.35 | 4239 | 1878.5 | **<0.001** |
| Prey identity× Log<sub>10</sub> predator body mass<sup>2</sup> | 8 | 19.05 | 4231 | 1859.5 | **0.0146** |

**Fig 2.**
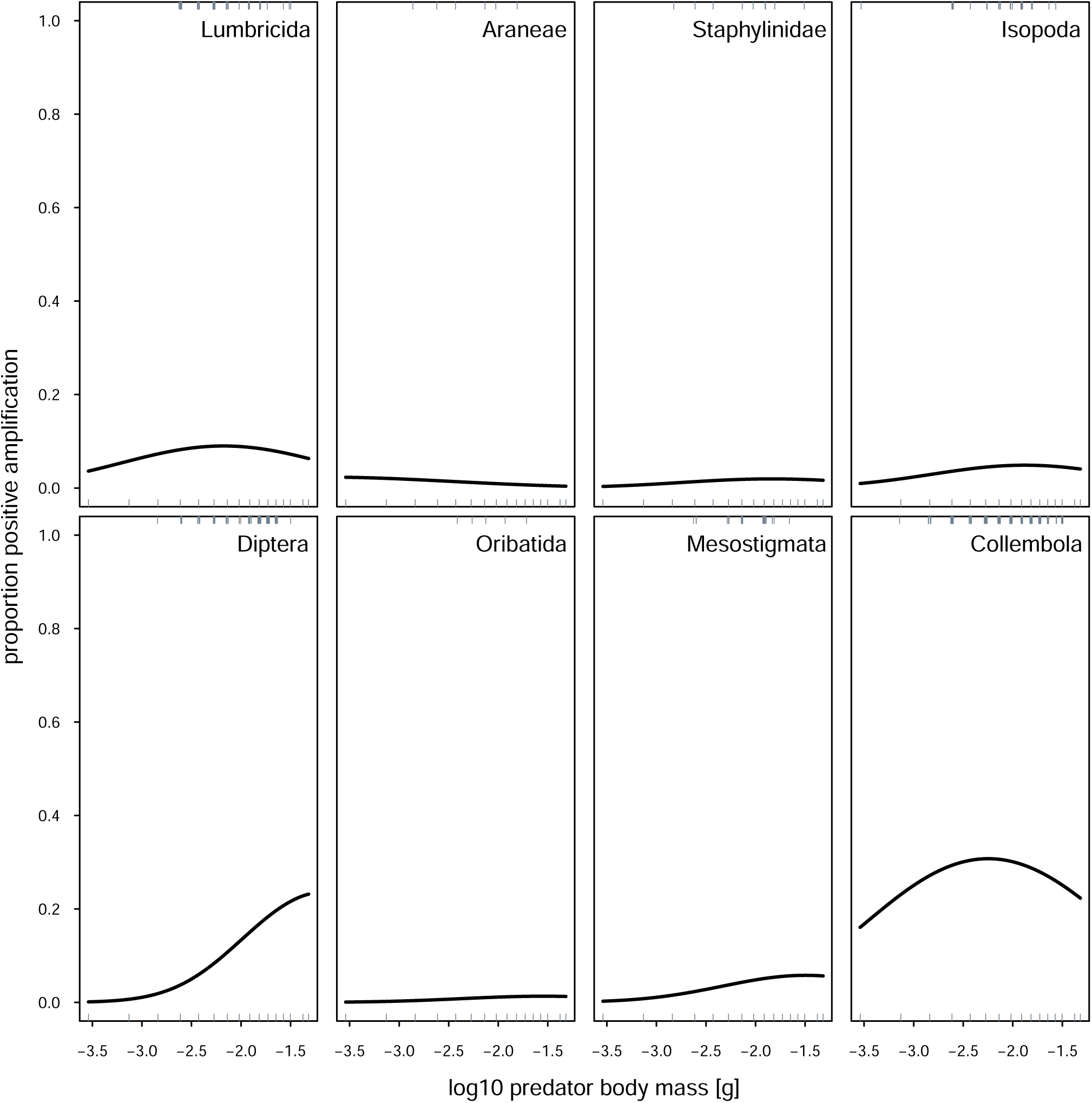
Body-size-dependent probability of positive prey-DNA detection of eight taxa in lithobiid centipedes (n= 532) sampled in autumn 2009 and spring 2010. Rugs on top and bottom of each diagram display single data points with values 1 or 0.

### Prey proportions according to functional response models

According to the functional response models, Collembola, Oribatida and Mesostigmata accounted for most of the diet of lithobiid and geophilomorph centipedes, showing a bimodul relationship with predator body mass (Fig. 3; Fig. S4, Supplementary Information). Diptera and Isopoda prey portions increased slightly at highest body masses, while other prey did not form part of the diet of the centipede predators.

**Fig. 3.**
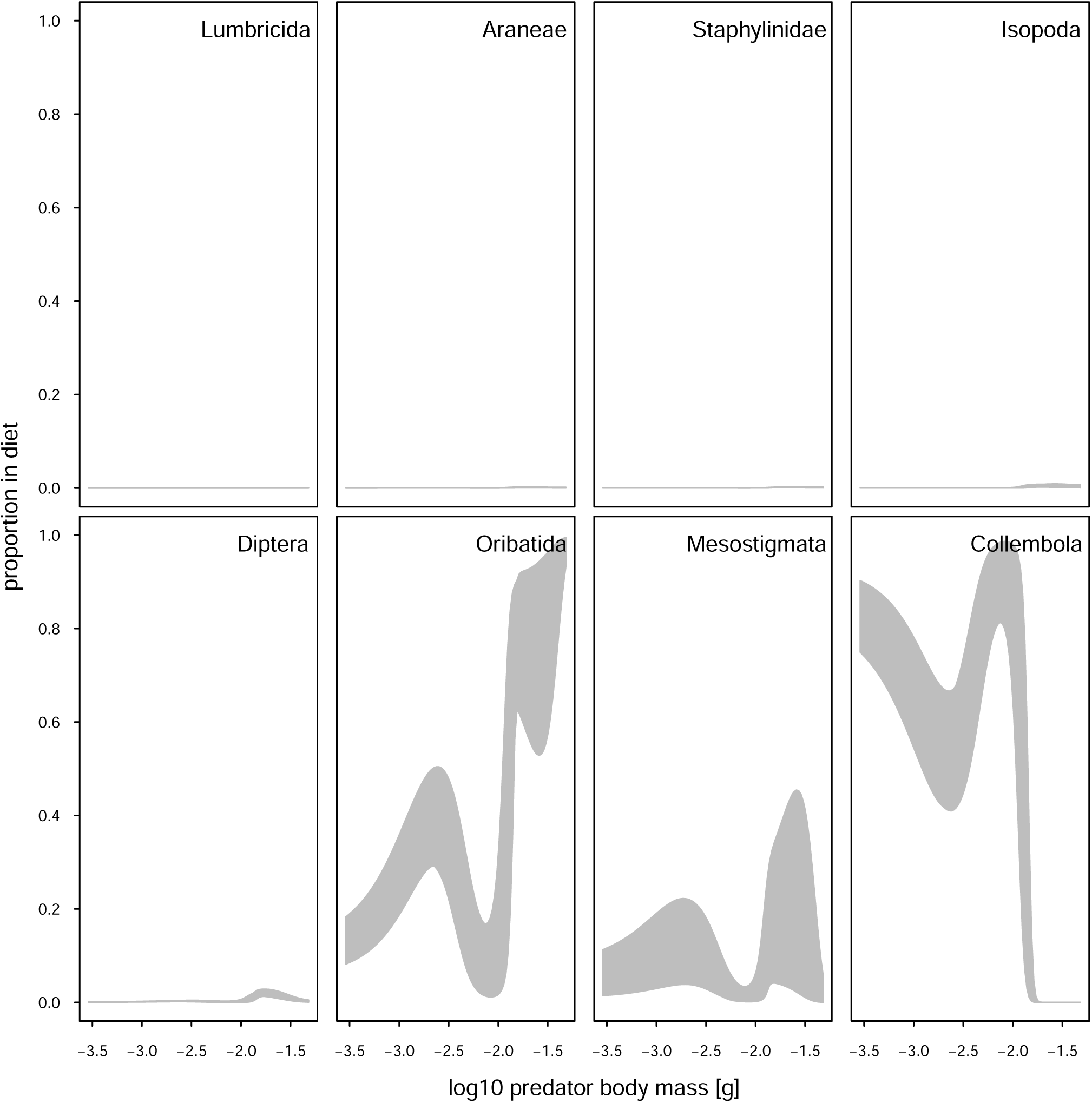
Body-size-dependent proportion of eight prey taxa in the diet of centipede predators as based on the functional response model using abundance and body-size data of invertebrates sampled in autumn 2009 and spring 2010. Upper and lower limit indicate highest and lowest diet proportion in the four forest sites.

### Comparison of functional response models with molecular gut content analysis

The relative proportion of a specific prey in the centipedes’ diet, as calculated by functional response models and the proportion of prey-DNA-positive centipedes, as calculated from the molecular gut content analysis significantly correlated for each of the prey group (Pearson correlation coefficient, *P* < 0.001; Fig. 4). While we found a positive correlation for the five prey groups Collembola, Diptera, Isopoda, Oribatida and Staphylinidae the other three prey groups had a negative relationship. In geophilomorph centipedes, only correlations with Lumbricidae, Staphylinidae and Collembola were significantly positive (*P* < 0.05), while Mesostigmata showed a significant negative correlation (*P* < 0.001). The other prey groups did not show any significant correlation.

**Fig 4.**
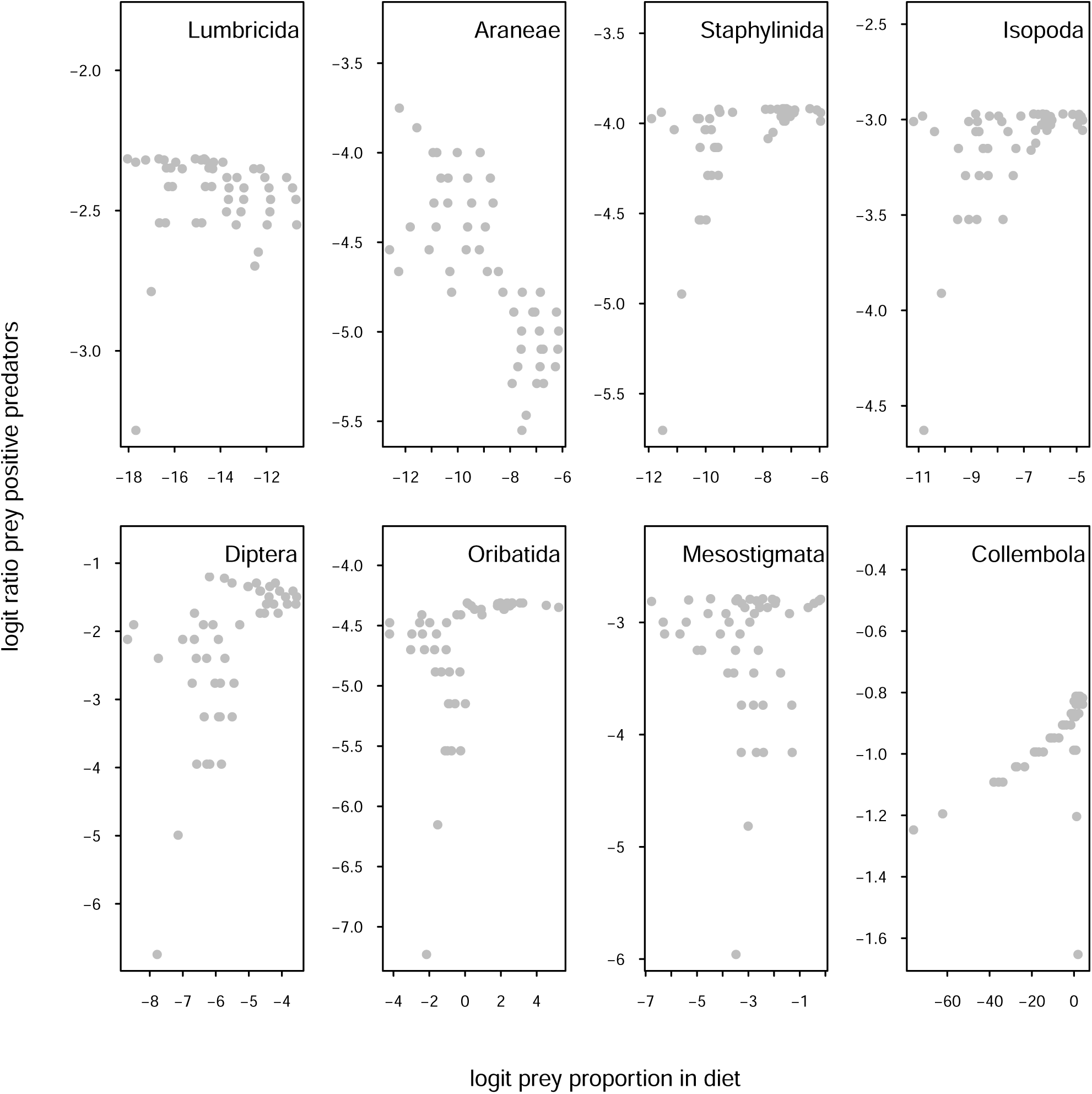
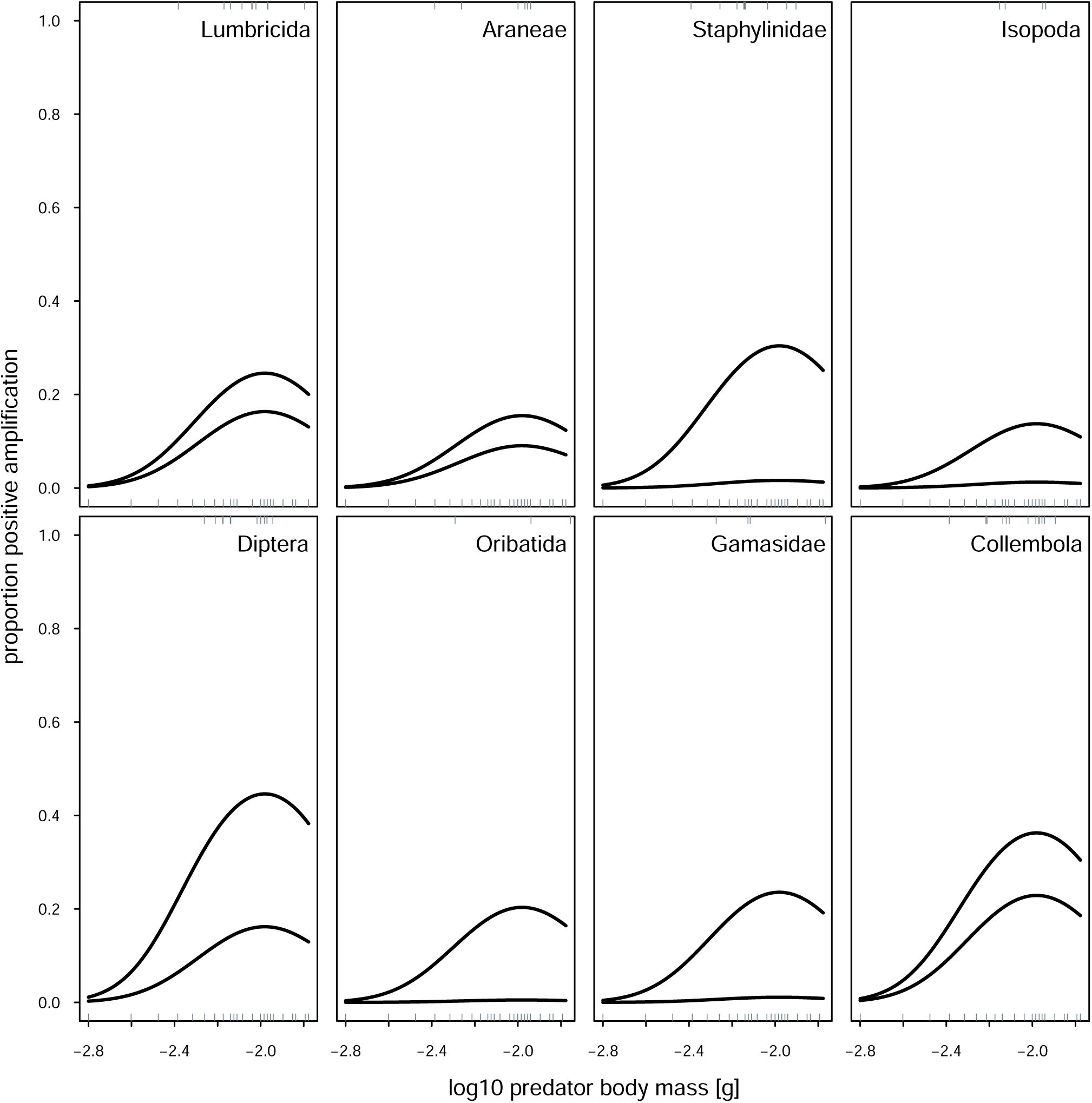
Pearson correlation coefficient between the relative proportion of prey in the centipede’s diet (as calculated by functional response models) and the proportion of prey-DNA-positive tested centipede *Lithobius* sp. (based on molecular gut content data) for each of the eight main prey groups.

## Discussion

The present study provides the first strong evidence that generalised allometric functional response models are an appropriate method to assess predator-prey interactions in complex systems, which include high levels of habitat structure, competitors and alternative prey. We tested if these models correctly predict relative feeding strength of generalist predators in a species- rich soil system by comparing with empirically quantified prey proportions in the diet of predators as indicated by molecular gut content analysis. Model and empirical data positively correlated in five of eight tested prey species, suggesting high explanatory power of the functional response models. Corroborating previous studies employing functional response models (Vucic-Pestic *et al.* 2010, Rall *et al.* 2011), we also empirically showed that ‘predator body size’ and ‘prey identity’ are two major drivers of prey capture in soil-dwelling predators.

The functional response models predicted high feeding rates of both lithobiid and geophilomorph centipedes on mesofaunal prey including Collembola, oribatid and mesostigmatid mites. A combination of high prey abundance, facilitating high encounter rates, and an optimal predator-prey body mass relationship allows the predator to forage on a maximum of prey individuals with a minimum of handling time, thereby reducing energetic costs (Aljetlawi *et al.* 2004, Brose *et al.* 2008, Vucic-Pestic *et al.* 2010). Results of the model used in this study allowing to track shifts from a hyperbolic (type-II) to a sigmoid (type-III) functional response suggest that with increasing predator body mass relative feeding rates follow a roller-coaster-pattern, peaking at the respective optimal body-mass ratios.

Feeding rates on other than mesofauna prey, however, were consistently low, only increasing slightly in large lithobiids and geophilomorph centipedes. As metabolism increases with body size, consumers require a higher energy uptake which is covered by the ingestion of more prey biomass, i.e. more small prey or larger prey individuals (Kalinkat *et al.* 2011). This is in line with earlier studies (Woodward & Hildrew 2002, Kalinkat *et al.* 2011) showing that with the increase in predator body mass prey preference shifts towards bigger prey while at the same time still being able to exploit small prey.

Results from the molecular gut content analysis corroborate the body-size dependent change in prey capture in the mathematical model. Centipedes exhibit unimodal feeding responses for 75% of the studied prey taxa, with large predator individuals more frequently feeding on more prey taxa than small predators. Analogous to the model, mesofauna taxa constitute the most important prey except for oribatid mites, which were detected in only 0.94% and 4.62% of the tested lithobiid and geophilomorph centipedes, respectively. While their high abundances and optimal body size suggest them to be ideal prey in the model, other traits, particularly their hard exoskeleton and toxic secretions seem to be effective defence traits, explaining why they were only rarely consumed (Peschel *et al.* 2006, Heethoff *et al.* 2011).

Collembola-DNA was detected in most centipedes, particularly medium-sized individuals. Collembola are abundant in virtually any terrestrial ecosystem and of high nutritional value thereby functioning as major prey for a wide range of predators in soil throughout the globe (Marcussen *et al.* 1999, Bilde *et al.* 2000, Oelbermann *et al.* 2008). Using a taxonomic-allometric model, Rall *et al.* (2011) calculated an optimal body mass ratio of 649 between the lithobiid centipede species *L. forficatus* and the Collembola species *Heteromurus nitidus.* In our study a similar ratio applied to *L. lanuginosus* and *P. armata,* the second and third most often detected Collembola prey species of lithobiid centipedes, respectively.

Lumbricidae, on the other hand, were a far more important prey than expected from the functional response model. Lumbricidae for long have been regarded as major prey of centipedes, in particular geophilomorph species (Lewis 1981), however, their low abundances and big size (as compared to mesofauna taxa) make them an unlikely prey in our allometric model. Using their poison claws, however, centipedes kill prey far below the optimal body-mass ratio (Eason 1964), and this resulted in underestimation of the importance of earthworms as prey of centipedes.

Interestingly, we found a strong increase in feeding on Diptera larvae with lithobiid body size, even stronger than predicted by the model. In combination with reduced feeding on other important prey, Collembola and Lumbricidae, this suggests prey switching towards this abundant prey of high nutritional value (Oelbermann & Scheu 2002). Prey switching has been reported in many studies (Hohberg & Traunspurger 2005, Petchey *et al.* 2008) and its frequency is increasing if predators become larger, presumably due to a combination of effects of habitat structure and optimal foraging processes (Murdoch & Oaten 1975, Kalinkat *et al.* 2013a) as described as follows:

Habitat structure modifies lithobiid feeding by allowing small prey such as Collembola but also small Lumbricidae, to take refuge from predation, forcing particularly large predator individuals to focus on more accessible prey dwelling in the upper litter layer (Günther *et al.* 2014).

Simultaneously, larger predators have higher energetic demands forcing them to hunt for larger prey, i.e. bigger individuals of species already feeding upon or a new, larger species. Higher energetic costs of killing, ingesting and digesting (i.e. ‘handling time’) prey, such as tipulid fly larvae or large earthworms are more easily balanced by the prey’s high nutritional value. However, the results suggest that to meet their nutritional and energetic demands, large lithobiid centipedes cannot be too selective in their prey choice: their spectrum still includes mesofauna prey and also encompasses intraguild prey, such as spiders and staphylinid beetles. These results confirm earlier studies showing that the prey spectrum of predators broadens with predator body size, suggesting that large predators exploit prey communities more efficiently (Cohen *et al.* 1993; Woodward & Hildrew 2002). On the other hand our findings argue against suggestions that at high density of extraguild prey intraguild predation is negligible (Halaj & Wise 2002, Eitzinger & Traugott 2011). Further, the results contradict findings that the role of intraguild predation is reduced in well-structured habitats providing refuge for intraguild prey (Finke & Denno 2002, Janssen *et al.* 2007).

## Conclusions

The present study, for the first time, investigated the impact of predator body size and prey abundance on predator consumption using two different approaches, functional response models and molecular gut content analysis. Both methods proved to be useful to study trophic interactions, the first one to analyse feeding strengths based on body size ratios and abundances, the latter to examine predator-prey interactions of individual predators on small scale. While these methods measure different parameters, i.e. feeding rate and prey DNA detection frequency, respectively, results of the present study suggest that they complement each other allowing to prove and extend theoretical predictions under natural settings. Therefore, combining these two techniques may ultimately allow uncovering the structure of food webs in particular those in opaque habitats colonized by minute animal species.

Combining functional responses with molecular gut content analyses and including predator-prey body size ratios we are able to explain the majority of feeding interactions in belowground systems. This emphasizes that allometric constraints override taxonomic constraints in structuring soil food webs. Further, in contrast to food webs in simply structured habitats, such as aquatic systems, prey abundance did not affect prey ingestion rates in this soil system, pointing to the importance of prey identity effects as driving factors. Therefore, for improving the effectiveness of allometric functional response models in predicting food web interactions in the field, additional traits of prey species, such as defence characteristics, have to be included.

## Author’s contributions

B.E. and B.C.R. conceived the ideas and designed methodology with contributions from M.T. and S.S.; B.E. collected the data, and B.E. and B.C.R. analysed the data; B.E. drafted the manuscript. All authors contributed to later drafts and gave final approval for publication.

## Acknowledgments

We thank the managers of the three Exploratories, Kirsten Reichel-Jung, Swen Renner, Katrin Hartwich, Sonja Gockel, Kerstin Wiesner, and Martin Gorke for their work in maintaining the plot and project infrastructure; Christiane Fischer and Simone Pfeiffer for giving support through the central office, Michael Owonibi for managing the central data base, and Markus Fischer, Eduard Linsenmair, Dominik Hessenmöller, Jens Nieschulze, Daniel Prati, Ingo Schöning, François Buscot, Ernst-Detlef Schulze, Wolfgang W. Weisser and the late Elisabeth Kalko for their role in setting up the Biodiversity Exploratories project. The work has been funded by the DFG Priority Program 1374 "Infrastructure-Biodiversity-Exploratories". Field work permits were issued by the responsible state environmental offices of Thüringen (according to § 72 Bbg NatSchG).

We particularly want to thank Olga Ferlian, Stephan Töppich and David Ott for their help with fieldwork, Bernhard Klarner for providing data on prey abundance and Christoph Digel and Amrei Binzer for help with statistics.

## Data accessibility

If the manuscript gets accepted, the authors will make data available on the Dryad Digital Repository (http://www.datadryad.org).

